# Multi-night naturalistic cortico-basal recordings reveal mechanisms of NREM slow wave suppression and spontaneous awakenings in Parkinson’s disease

**DOI:** 10.1101/2023.06.23.546302

**Authors:** Md Fahim Anjum, Clay Smyth, Derk-Jan Dijk, Philip Starr, Timothy Denison, Simon Little

## Abstract

**Background:** Sleep disturbance is a prevalent and highly disabling comorbidity in individuals with Parkinson’s disease (PD) that leads to worsening of daytime symptoms, accelerated disease progression and reduced quality of life.

**Objectives:** We aimed to investigate changes in sleep neurophysiology in PD particularly during non-rapid eye movement (NREM) sleep, both in the presence and absence of deep brain stimulation (DBS).

**Methods:** Multi-night (n=58) intracranial recordings were performed at-home, from chronic electrocorticography and subcortical electrodes, with sensing-enabled DBS pulse generators, paired with portable polysomnography. Four people with PD and one person with cervical dystonia were evaluated to determine the neural structures, signals and connections modulated during NREM sleep and prior to spontaneous awakenings. Recordings were performed both ON and OFF DBS in the presence of conventional dopaminergic replacement medications.

**Results:** We demonstrate an increase in cortico-basal slow wave activity in delta (1-4 Hz) and a decrease in beta (13-31 Hz) during NREM (N2 and N3) versus wakefulness in PD. Cortical-subcortical coherence was also found to be higher in the delta range and lower in the beta range during NREM versus wakefulness. DBS stimulation resulted in a further elevation in cortical delta and a decrease in alpha (8-13 Hz) and low beta (13-15 Hz) power compared to the OFF stimulation state. During NREM sleep, we observed a strong inverse interaction between subcortical beta and cortical slow wave activity and found that subcortical beta increases prior to spontaneous awakenings.

**Conclusions:** Chronic, multi-night recordings in PD reveal opposing sleep stage specific modulations of cortico-basal slow wave activity in delta and subcortical beta power and connectivity in NREM, effects that are enhanced in the presence of DBS. Within NREM specifically, subcortical beta and cortical delta are strongly inversely correlated and subcortical beta power is found to increase prior to and predict spontaneous awakenings. We find that DBS therapy appears to improve sleep in PD partially through direct modulation of cortico-basal beta and delta oscillations. Our findings help elucidate a contributory mechanism responsible for sleep disturbances in PD and highlight potential biomarkers for future precision neuromodulation therapies targeting sleep and spontaneous awakenings.

## INTRODUCTION

Sleep disruption is one of the most prevalent non-motor symptoms of Parkinson’s disease (PD) with up to 90% of PD patients experiencing sleep dysfunction^1^ and 60% having multiple sleep disturbance symptoms^1,2^. Changes in sleep patterns often predate classical neurological symptoms in PD and correlate with rates of progression and disease severity^3^. Sleep dysfunction in PD has a negative impact on daytime mood, cognition, fatigue, and other co-morbidities^4–8^, with non-motor and sleep symptoms being a greater determinant of quality of life than classical motor symptoms^9–11^. Therefore, understanding the neurophysiology of sleep disturbances in PD may potentially result in new principled therapies directed towards better sleep quality, mitigation of daytime symptoms and improved patients’ quality of life.

Sleep architecture in humans is broadly defined by physiologically distinct stages of rapid eye movement (REM) and non-REM (NREM) sleep. NREM sleep is further characterized by rhythmic low frequency electroencephalography (EEG) activity in the delta (0-4 Hz) and theta (4-7 Hz) ranges, increased parasympathetic activity and limited dreaming. There are currently three formally defined sub-stages of NREM: N1 (light sleep), N2 (appearance of K complexes and sleep spindles) and deep N3 (characterized by slow delta waves)^13^. Sleep dysfunction in PD manifests as parasomnias, fragmented sleep and disrupted sleep patterns, including notable reductions in both REM and NREM sleep^11^. In particular, reductions in NREM sleep slow wave activity in the delta range (< 4Hz) are associated with worsening of daytime motor symptoms and accelerated disease progression in PD^3,14,15^.

During wakefulness, beta oscillations (13-30 Hz) are the hallmark oscillatory signature of PD and correlate with daytime motor symptoms^16^. Recent studies with non-human primates (NHPs) during sleep have shown that subcortical beta activity is also associated with a decrease in cortical delta activity and suggested a role for subcortical beta in spontaneous awakenings in PD^17^. The presence of beta oscillations has been detected during sleep in PD patients^18–22^. However, to date, human studies have not yet investigated mechanistic interactions between subcortical beta and cortical sleep physiology (inc. slow waves) nor spontaneous awakenings and previous studies have been completed in the absence of deep brain stimulation (DBS). Understanding the real-world contribution of the cortico-basal ganglia circuit to sleep dysfunction in PD and its interaction with DBS, has to date been limited by an inability to chronically record the intracranial activities overnight, at high resolution. This challenge has been solved by the advent of a new generation of sensing-enabled DBS devices that can stream neural data remotely from patients’ own homes^23^. A better understanding of cortico-basal activities during sleep has the potential to reveal underlying mechanisms of sleep dysfunction in PD and could contribute to improved sleep therapies including sleep-targeted adaptive deep brain stimulation (aDBS).

In this study, we recruited four patients diagnosed with PD and one comparison patient with cervical dystonia, all with chronically implanted intracranial electrodes capable of sensing sensorimotor cortical and basal ganglia (STN/GPi) field potentials (FPs). We conducted overnight, at-home, intracranial cortical and subcortical recordings paired with portable polysomnography over multiple nights (n= 58) in the presence and absence of DBS stimulation. We demonstrate significant interactions between subcortical beta oscillations and cortical slow wave activity in the delta band during NREM, an effect modulated by DBS, and also show that subcortical beta significantly increases prior to spontaneous awakenings.

## RESULTS

Four people (Table 1) with PD (x2 people with bilateral STN + sensorimotor cortical ECoG and x2 people with bilateral GPi electrodes + sensorimotor cortical ECoG) and one person with cervical dystonia (bilateral GPi electrodes + sensorimotor cortical ECOG), successfully initiated recordings from intracranial cortico-basal and external portable polysomnography (Dreem2) over 58 nights (54 ON and 4 OFF stimulation nights), remotely in their own homes. Intracranial and extracranial recordings were synchronized and artifacts were removed (Fig 1E; Supplementary Fig. 3), resulting in interpretable cortical and subcortical recordings, even in the presence of DBS. A total 415 hours of sleep were recorded across all participants (Supplementary Table 1; Supplementary Fig. 1). PD subjects slept on average 7.25±0.18 hours per night during the extended multi-night ON stimulation (n=45; duration in minutes: N1 = 34.26±1.34; N2 = 164.72±7.92; N3 = 90.06±10.53; REM = 97.64±6.54; Wake after sleep = 48.18±4.71). In the separate two night consecutive ON versus OFF DBS comparison recording nights, all four PD subjects showed an increase in time in deep NREM (N3) and REM in the ON stimulation compared to the corresponding OFF stimulation nights (Supplementary Table 1 & Fig. 2). Power spectral density plots from intracranial electrodes (Fig. 1G and Supplementary Fig. 4) demonstrated expected classical changes in canonical frequency bands in NREM and REM sleep stages, supporting dissociation of different sleep stages using our portable PSG device sleep staging.

**Table 1:** Participant demographics.

| Subject ID | PD2 | PD3 | PD7 | PD9 | Dystonia |
| --- | --- | --- | --- | --- | --- |
| Age | 58 | 66 | 40 | 48 | 65 |
| Gender | M | M | M | M | M |
| Diagnosis | PD | PD | PD | PD | Dystonia |
| Dx | 11 | 13 | 9 | 13 | 30 |
| RCS Target | STN | GPI | STN | GPI | GPI |
| Pulse Width | 60 | 60 | 60 | 90 | 60 |
| Stimulation Frequency | 130.2 | 178.6 | 130 | 150 | 130.2 |
| L contact | C+2- | C+1- | C+2- | C+2- | C+1- |
| R contact | C+1- | C+1- | C+2- | C+2- | C+2- |
| Medication | A-HCL 100mg (3 times daily)<br>C-Ldopa 25-100 mg IR (5 times daily) | C-Ldopa 25-100mg CR (1-2 tabs at bedtime) and 25-100mg IR (3 times daily) | C-Ldopa 25-100mg (1 time daily) Rasagaline (Azilect) 1mg (1 time daily) | Rytary 195mg (3 times daily) | NA |
| UPDRS-III (OFF) | 49 | 66 | 41 | 39 | NA |
| UPDRS-III (ON) | 5 | 24 | 14 | 16 | NA |
| UPDRS 1.7 | No sleep symptoms | Slight sleep symptoms | Slight sleep symptoms | Mild sleep symptoms | NA |
| UPDRS 1.8 | No daytime sleepiness | Mild daytime sleepiness | Moderate daytime sleepiness | Mild daytime sleepiness | NA |
| Sleep diagnosis | No sleep sx/conditions | Nocturia, RBD | Daytime sleepiness | OSA, Insomnia | Restless Leg Syndrome |
| Neuropsych report (pre-op) | No reported sleep disorder or conditions | Mild sleep difficulties, with nocturia and RBD | Day time sleepiness (strongest 4-5pm), usually sleeps late and sleeps very little overnight | Had difficulties sleeping before PD. Occasionally couldn't fall asleep at night. | Good sleep. No movements/dystonia at night. Restless Leg Syndrome at night. |

UPDRS scores were pre-op, Dx = Disease duration (Years), RBD = REM sleep behaviour disorder, Obstructive sleep apnea = OSA, PD = Parkinson's disease, C-Ldopa = Carbidopa-Levodopa (Sinemet), A-HCL = Amantadine HCL (Symmetrel). None of them suffered from Dementia.

**Figure 1.**
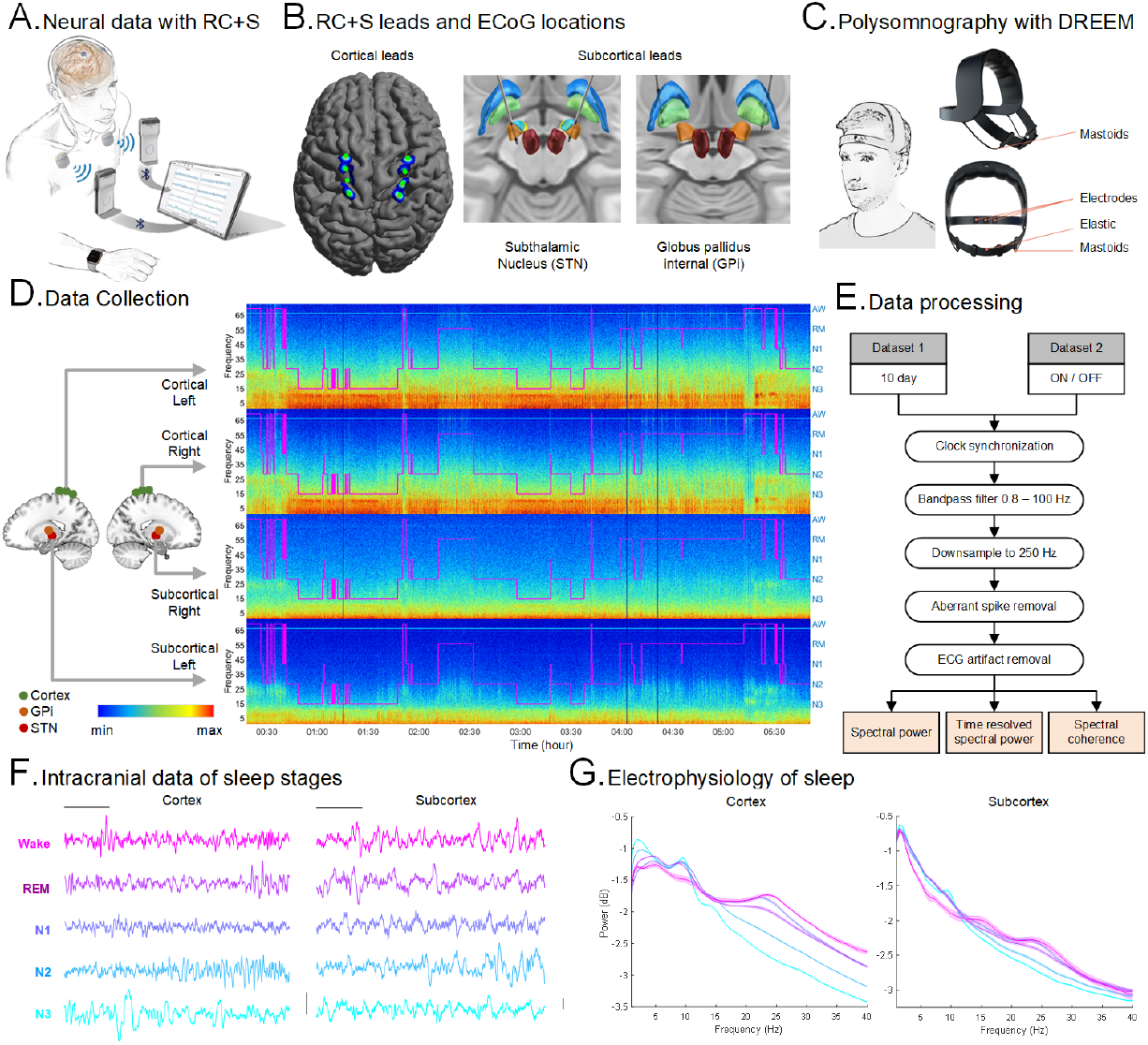
Methodology, data collection and analysis procedures: **(A)** Schematic of the RC+S system setup for recording intracranial cortical Field Potentials (FP) in participants (Adapted from Gilron et al. 2021^23^). **(B)** Illustrations of the placement of RC+S sensing depth electrodes in subcortex for both STN and GPi (*right*) and cortical ECoG locations (*left*). Example data from PD2 and PD3 participants. **(C)** Schematic of the Dreem2, portable headband for recording in-home polysomnography over night (adapted from Debellemaniere et al. 2018^36^). **(D)** Illustration of a single night of sleep in a PD patient (DBS ON) with hypnogram (purple) showing sleep stages (AW: awake; RM: REM; [N1, N2, N3]: NREM) and cortical (top 2 panels) and subcortical (bottom 2 panels) spectrogram panels from both hemispheres showing multi-frequency changes across sleep stages where the x-axis is time in hours and y-axis is frequency (Hz). FP was recorded bilaterally from cortical and subcortical regions. **(E)** Flowchart of data analysis and preprocessing procedures for 10 day sleep dataset (n=5) and ON/OFF dataset (n=4). **(F)** Representative traces of the RC+S FP time series in all sleep stages from cortex (*left column*) and subcortex (*right column* ;Subthalamic Nucleus). Columns share scale bars and rows share color legends (Wake, REM, N1, N2 and N3). Data from one PD participant with ON stimulation from the left hemisphere. **(G)** Comparisons of spectral powers of intracranial FPs among sleep stages in cortex (*left*) and subcortex (*right*) for a single subject, DBS ON. Shaded error bars indicate standard error. Shares color legend with panel F.

**Figure 2:**
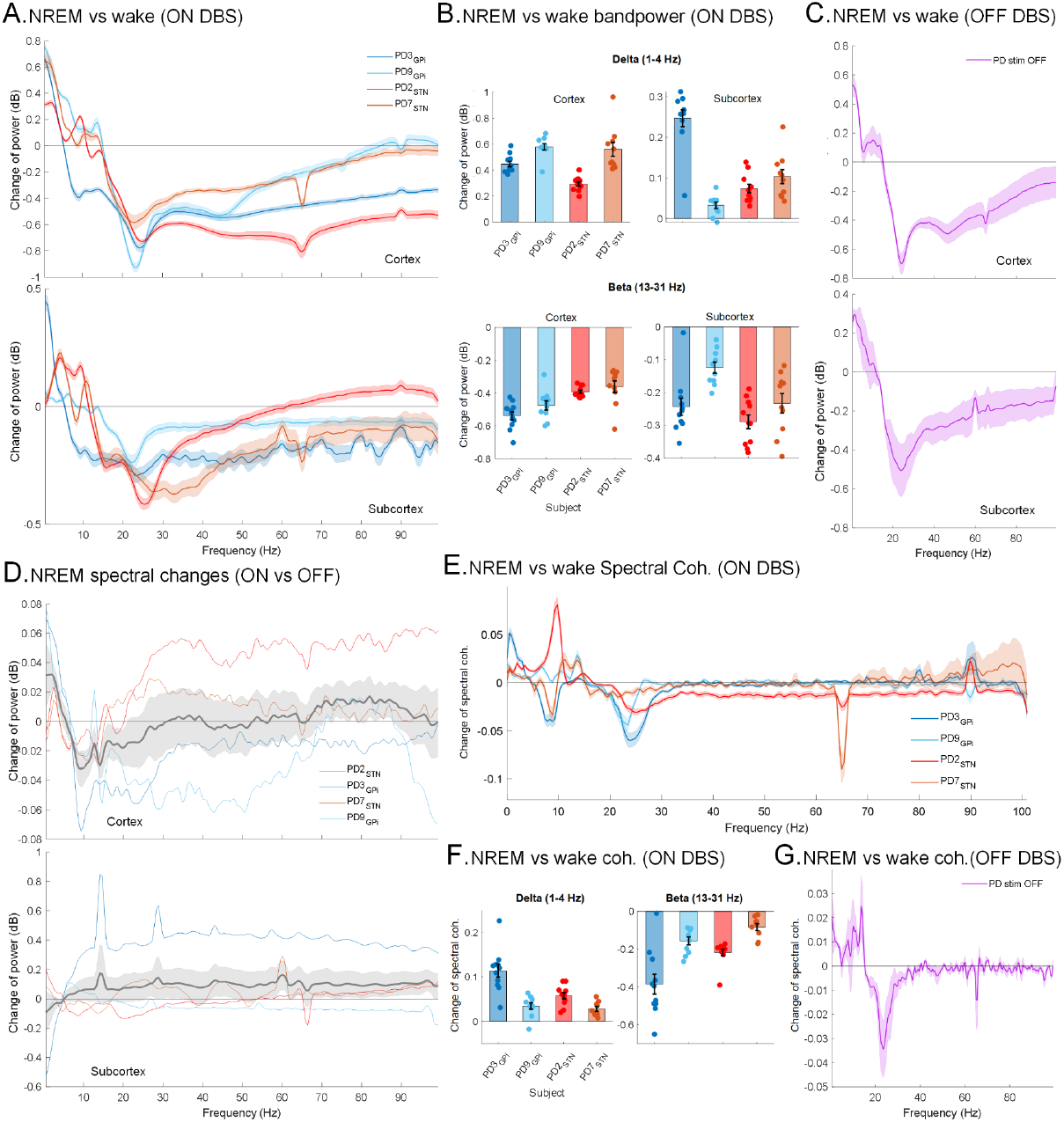
Spectral changes in NREM. Dynamic changes in power spectra and functional connectivity between cortical and subcortical regions during NREM sleep: **(A)** Power Spectrum changes (mean ± SEM) during NREM (N2 and N3) sleep with wake stage as baseline in low frequency range (1-50 Hz) for all PD participants (n=4) during ON stimulation in cortical (*top*) and subcortical (*bottom*) areas. y-axis shows the difference in power spectra between NREM and wake stage in decibel (dB). Thick lines show mean and shaded areas show standard errors (SEM). **(B)** Power in delta (1-4 Hz) increases while beta (13-31 Hz) band power decreases during NREM sleep compared to wake during ON stimulation in cortical (*top*) and subcortical (*bottom*) areas. Each bar shows the difference in spectral power for one participant averaged across multiple nights and each data point shows the average difference in spectral power across one night with data pooled from both hemispheres. **(C)** During OFF stimulation conditions, delta power increases while beta power decreases in NREM compared to the wake stage in PD participants (n=4) in cortical (*top*) and subcortical (*bottom*) areas. Thick lines show means and shaded areas show standard errors. **(D)** Difference in cortical spectral power between ON and OFF stimulation conditions in 4 participants with PD in NREM sleep stages (*top*), showing increased delta (1-4 Hz) and decreased low-alpha and low-beta activities (8-15 Hz) while ON stimulation. Each colored line shows spectral change for one participant, thick line shows average across the participants with shaded area as SEM. The spectral power in subcortical regions didn’t show any statistically significant difference (*bottom*). The x-axis is frequency (Hz) and the y-axis is difference in power (ON-OFF). **(E)** Changes in cortical-subcortical spectral coherence (mean ± SEM) during NREM (N2 and N3) sleep with wake stage as baseline for all participants (n=5) during ON stimulation. y-axis shows the difference in spectral coherence between NREM and wake stage. Horizontal back line at 0 represents wake stage baseline. **(F)** Total difference in spectral coherence in delta (1-4 Hz, *left*) and beta (13-31 Hz, *right*) during NREM sleep compared to wake during ON stimulation. Each bar shows difference in spectral coherence for one participant averaged across multiple nights and each point shows average difference in spectral coherence across one night with data pooled from both hemispheres. **(G)** During OFF stimulation conditions, delta coherence increases while beta coherence decreases in NREM compared to the wake stage in PD participants (n=4). Data from both hemispheres were pooled for all panels.

**Figure 3.**
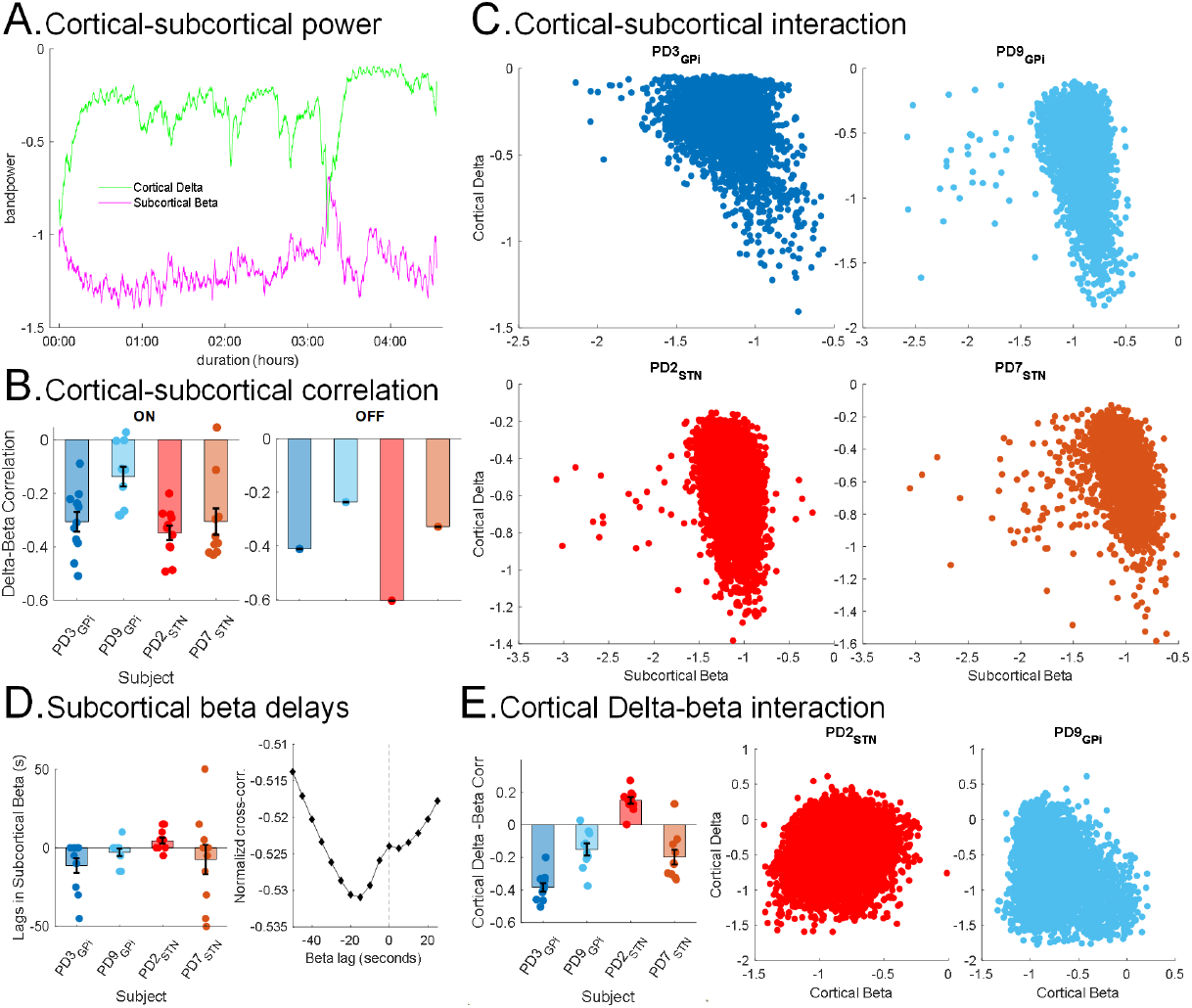
Inverse relationship between subcortical beta and cortical delta activities. **(A)** Example of subcortical beta (*purple*) and cortical delta (*green*) power in a single night from one PD participant (PD3) during ON stimulation depicting the inverse relationship in temporal domain. The delta and beta powers were smoothed with a 20-point gaussian kernel. **(B)** Average Spearman’s rho correlation between subcortical beta power and cortical delta power for all 4 PD participants across multiple nights in ON (*left*) and in OFF (*right*) stimulation. Each bar shows average correlation for one participant and each point shows correlation across one night with data pooled from both hemispheres. **(C)** Scatter plots depicting the correlation between subcortical FP beta (13-31 Hz) power and cortical FP delta (1-4 Hz) power during NREM sleep in 4 PD participants during ON stimulation; STN (*brown and red*), and GPi (*blue, light blue*). Each point represents data from one 5-s NREM sleep epoch. Each plot is data from one night pooled from both hemispheres for one participant. **(D)** Normalized cross-correlation between subcortical beta power and cortical delta power showing the subcortical beta preceding cortical delta activities in PD participants during NREM with ON stimulation. The bar plot (*left*) shows lags in subcortical beta with cortical delta as reference. Each bar shows average lag for one participant and each point shows lag across one night with data pooled from both hemispheres. Example of cross correlation showing the lag in subcortical beta as a function of time (*right*) in one night from PD2 during ON stimulation. The vertical dashed line shows zero-lag. **(E)** Interactions between cortical delta and cortical beta activities, examined as a control for cortical delta - subcortical beta. The bar plot (*left*) shows average Spearman’s rho correlation between cortical delta and beta power for all 4 PD participants across multiple nights, ON stimulation. Each bar shows average correlation for one participant and each point shows correlation across one night with data pooled from both hemispheres. The scatter plots show cortical delta and beta power in 4 PD participants during ON stimulation for two representative PD participants. Each point represents data from one 5-s NREM sleep epoch.

**Figure 4:**
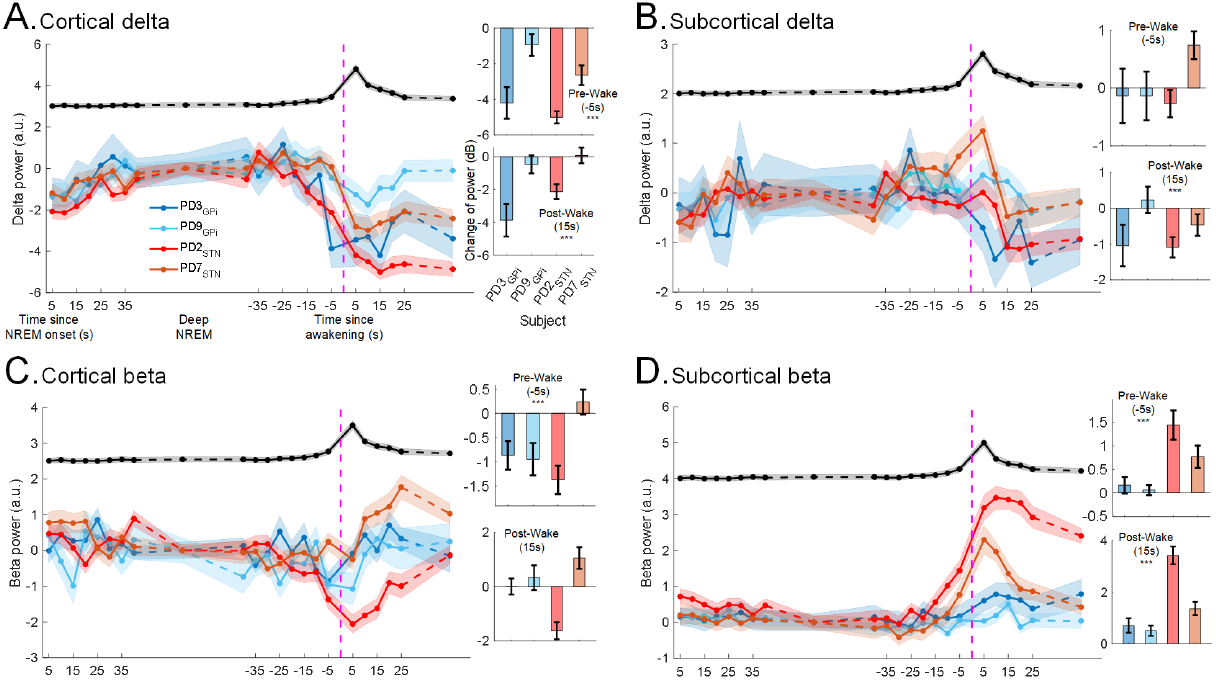
Changes in spectral power before spontaneous awakenings. Subcortical beta increases and cortical delta decreases before spontaneous awakening. **(A)** Cortical delta (1 - 4 Hz) power during NREM to wake sleep transition episodes for all PD participants (n=4; mean ± SEM) during ON stimulation (*left*). Each data point is the average for 5-s data epochs and shadings represent SEMs for NREM to wake transitions across the recording nights for one participant. Data were pooled from both hemispheres. The vertical purple dashed line shows awakening time. x-axis (*on the left*) shows time in seconds since NREM sleep onset and time since awakening (*middle*, around vertical dashed line). The black line on top shows the norm of RC+S accelerometry data (mean ± SEM) for all NREM to wake transitions across all nights for all participants highlighting the awakening time of the episodes. The bar plots show change in cortical delta power during pre-awakening (5-s before the wake event, *top*) and post-awakening (15-s after the wake event, *bottom*) compared to the average delta power in deep NREM (average over NREM data after 40-s from NREM onset and 40-s before awakening; SWS). Each bar shows average change of power for one participant and each point shows change of power across all NREM to wake transitions in one night with data pooled from both hemispheres. Cortical delta power gradually increases as sleep deepens and decreases steadily before awakening. The average post-awakening (15s) and pre-awakening delta powers (−5s) are lower than those during SWS. The average post-awakening (15s) cortical delta power is lower than the pre-awakening delta power (−5s). **(B)** Same as A, for subcortical delta power showing no significant trend across participants or recording sites. **(C)** Same as A, for cortical beta power showing no significant trend across participants or recording sites for pre and post awakenings. **(D)** Same as A, only for subcortical beta power illustrating spontaneous rise in beta power before awakenings. The average post-awakening (15s) and pre-awakening beta power in STN (−5s) are higher than those during SWS.

### Spectral power changes in NREM

We investigated spectral changes in intracranial FP activities during NREM (N2 and N3) with wakefulness as a baseline with an a priori focus on cortico-basal delta and beta^24^. Power spectrum analyses and Linear Mixed Effect (LME) models for average overnight band powers with a fixed effect for sleep stage (NREM vs Wake; accounting for multiple nights within participants) and a random effect for subjects (n=5) showed an increase in average delta power (1 - 4 Hz; cortex: *β* = 0.42, 95%CI= [0.36, 0.47], p-value = 3.7e-33; subcortex: *β* = 0.1, 95%CI= [0.07, 0.13], p-value = 3.3e-12; n=105; CI=confidence interval) and decrease in beta (13 - 31 Hz; cortex: *β* = -0.4, 95%CI= [-0.44, -0.37], p-value = 1.5e-41; subcortex: *β* = -0.2, 95%CI= [-0.22, -0.17], p-value = 1.4e-25) power both in cortical and subcortical regions in NREM sleep compared to wakefulness (Fig. 2A-B; multi-night ON stimulation). These spectral changes in NREM compared to wake were also seen during the single night of OFF DBS sleep recordings in both cortical and subcortical regions of all four PD participants (Fig. 2C).

A direct comparison of our PD (n=4) vs Dystonia (n=1) analyses revealed that subcortical beta power was lower in the dystonia patients than all four of the PD patients during NREM sleep (LME model for PD vs Dystonia fixed effect: *β* = 0.18; 95%CI= [0.09, 0.27]; p-value = 0.0001). Further, band power changes between NREM sleep and wake condition in the dystonia participant were smaller compared to the PD patients (Fig. 2B). LME models demonstrated statistically significant fixed effects of disease state (PD vs Dystonia) on the changes of band power between NREM and wake stage in cortex (delta: *β* = 0.24, 95%CI= [0.14, 0.35], p-value = 1.3e-5; beta: *β* = -0.21; p-value = 3.4e-7) and subcortex (delta: *β* = 0.08, 95%CI= [0.013, 0.14], p-value = 0.02; beta: *β* = -0.13, 95%CI= [-0.2, -0.07], p-value = 0.0001).

We also investigated how these FP activities alter with stimulation and compared power spectrums between ON and OFF stimulation conditions in our PD cohort (n=4). Spectral power comparisons revealed a relative further increase in delta and a further decrease in alpha and low beta activities in cortical FP during NREM sleep in the ON versus OFF DBS conditions (Fig. 2D). LME models with subjects as random effects revealed that stimulation resulted in a further increased cortical delta (1-4 Hz; *β* = 0.026, 95%CI= [0.003, 0.05], p-value = 0.03) and decreased cortical alpha (8-13 Hz; *β* = -0.0297, 95%CI= [-0.05, -0.003], p-value = 0.03), low beta (13-15 Hz; *β* = -0.026, 95%CI= [-0.042, -0.01], p-value = 0.006) in the ON versus OFF condition, for NREM versus wakefulness. No significant changes in subcortical FP during NREM were observed in ON vs OFF power spectrum comparisons, however, changes in subcortical baseline power levels in the ON versus OFF DBS state may have obscured any underlying changes. Overall, these data reveal that DBS results in relatively higher cortical delta activity and reduced alpha and low beta activities in NREM sleep.

### Changes in functional connectivity in NREM

We next explored NREM-related changes in the functional connectivities between cortical and subcortical regions to investigate sleep-related changes in cortico-basal ganglia circuitry in PD. For this, we compared the spectral coherence in cortical and subcortical FP activities between NREM sleep and wakefulness. In all participants, LME models investigating spectral coherence with a fixed effect of sleep stage (NREM vs Wake) revealed that the total difference in spectral coherence in delta increases (*β* = 0.05, 95%CI = [0.04, 0.06], p-value = 5e-11; n=104) while beta decreases (*β* = -0.18; 95%CI = [-0.23, -0.14], p-value = 5.5e-13) during NREM sleep compared to wake in ON stimulation (Fig. 2E-F). An increase in delta coherence and a decrease in beta coherence during NREM were also observed in the PD participants during their single-night recordings OFF stimulation (Fig. 2G). PD vs Dystonia comparison also showed that cortio-basal delta / beta coherence changes in NREM versus wakefulness were smaller in the dystonia participants compared to the PD participants (LME model with PD/Dystonia condition as a fixed effect; delta coherence: *β* = 0.05, 95%CI= [0.02, 0.08], p-value = 0.0008; beta coherence: *β* = -0.21, 95%CI= [-0.31, -0.11], p-value = 0.0001).

In our ON vs OFF DBS analysis (PD participants only; n=4), we also noted a statistically significant further decrease in low beta (13 - 15 Hz) coherence during ON stimulation compared to OFF (LME model with ON/OFF condition as fixed and subjects as random effects; *β* = -0.012, 95%CI= [-0.021, -0.003], p-value = 0.015). Collectively, these data demonstrate that functional connectivity between cortical and subcortical structures is modulated during NREM sleep versus wakefulness. There is an increase in delta coherence and a decrease in beta coherence in PD in both ON and OFF stimulation conditions, effects that are enhanced in the DBS ON condition.

### Interaction between cortical delta and subcortical beta activity

Spectral power and functional connectivity analyses above revealed opposing changes in delta and beta FP activities in NREM sleep versus wakefulness. To further examine for a direct relationship between these two rhythms, we investigated the interactions between cortical delta and subcortical beta FP activities specifically *within* NREM (N2 + N3) on short time scales. Here, we observed an inverse relationship between cortical delta power and subcortical beta power during NREM sleep (Fig. 3A). To quantify this relationship, we first used a standard correlation analysis which revealed a negative correlation between subcortical beta and cortical delta FP power (5 s epochs) in all PD participants during NREM in both ON and OFF stimulation conditions (Fig. 3B-C). LME modeling using band powers of NREM epochs from all participants (Cervical dystonia and PD participants; accounting for the dependency between left and right hemispheres and multiple nights within patients; n=232,064) showed an overall negative fixed effect of subcortical beta power on cortical delta power (*β* = -0.24, 95%CI: [-0.28, -0.2], p-value = 3.9e-30). Additionally, we found a fixed effect of PD vs Dystonia state (*β* = 0.06, 95%CI: [0.01, 0.11], p-value = 0.02) during NREM sleep ON stimulation, demonstrating that this effect was greater in the PD patients that our dystonia comparison subject. Negative fixed effect of subcortical beta power on cortical delta power was also obtained through LME model in PD participants during the OFF stimulation condition (*β* = -0.38, 95%CI: [-0.44, -0.32], p-value = 1.9e-32; n=17,518). These results demonstrate that there is an inverse relationship between subcortical beta and cortical delta FP power within NREM sleep in PD both during ON and OFF stimulation conditions and that this effect is significantly stronger than in our comparison dystonia patient.

Next, we utilized cross-correlation analyses to determine whether subcortical beta was leading or lagging cortical delta changes. We observed that the subcortical beta increase was leading the cortical delta decrease in 3 out of the 4 PD participants during NREM sleep (Fig. 3D; average lag over multiple nights ON DBS; PD2: 4.5s, n=11; PD3: -11.4s, n=11, PD7: -7.5s, n=10, PD9: -3s, n=10). Finally, as a control analysis to rule out a prosaic inverse relationship between cortico-basal circuit delta and beta, simply reflecting depth of NREM sleep, we also correlated cortical delta and cortical beta power from the same region. If the inverse relationship between cortico-basal delta and beta was simply a function of sleep stage depth, we would expect the inverse relationship between cortical delta and cortical beta to be strongly inverted. Unlike correlations between subcortical beta and cortical delta, which were negative for all PD participants, cortical delta and beta showed a weaker negative correlation in only 3 PD participants and a positive correlation in one PD participant (Fig. 3E) as well as in the Dystonia participant during NREM (ON stimulation). LME analysis did show an overall negative fixed effect of cortical beta power on cortical delta power (*β* = -0.21, 95% CI: [-0.32, -0.1], p-value = 0.0002; n=232,064) during NREM sleep but did not show any fixed group effect of PD/Dystonia state (*β* = -0.02, 95% CI: [-0.06, 0.004], p-value = 0.1), indicating that at the cortical level, there was no evidence of any difference between the PD/dystonia conditions, in contrast to subcortically. Additionally, in direct model comparison, the LME model for cortical delta with a fixed effect of subcortical beta showed a statistically significant improvement over the model of cortical delta with a fixed effect of cortical beta (simulated likelihood ratio test with 100 replications; p-value = 0.01). This demonstrates that subcortical beta had a stronger effect on cortical delta compared to the relationship between cortical beta activity and cortical delta activity supporting that this subcortical beta - cortical delta effect is greater than any effect of sleep stage depth. Additionally, our data showed that subcortical beta-cortical delta effect was relatively specific to PD.

### Changes in spectral power before spontaneous awakenings

To better understand FP activities at a finer time resolution and investigate the dynamics of intracranial FP that lead to awakenings, we analyzed the change in spectral powers in delta and beta during NREM to spontaneous wake transitions. There were on average 26.7±1.5 awakening events per night, with a total duration of 52.7±4.5 minutes for each participant ON stimulation. During NREM, cortical delta power gradually increases as sleep deepens and decreases before awakening (Fig. 4A) in all participants (multi-night ON stimulation dataset). The average cortical delta power in pre-awakening (−5s) and post-awakening (+15s) periods were both lower compared to the average spectral power found in deep NREM stage, (Fig. 4A; pre-wake: *β* = -1.3, 95%CI: [-1.9, -0.6], p-value = 7.4e-5; n=446; post-wake: *β* = -2.9, 95%CI: [-3.5, -2.3], p-value = 7.7e-20) with the post-awakening cortical delta power being lower than the pre-awakening (Fig. 4A; *β* = -2.9 vs -1.3) and no fixed effects of PD/dystonia condition (pre-wake: p-value = 0.5; post-wake: p-value = 0.06). This suggests that changes in cortical delta are not PD-specific, but rather a general feature of changes in neurophysiology in NREM sleep versus wakefulness. The subcortical delta (pre-wake: p-value = 0.6; post-wake: p-value = 0.002) and cortical beta power (pre-wake: p-value = 0.00004; post-wake: p-value = 0.98) did not demonstrate consistent pre and post wake changes which were significant (Fig. 4B-C) across participants, around the time of spontaneous awakenings. However, the subcortical beta power demonstrated a rise before awakenings which was sustained after awakening (Fig. 4D; pre-wake: *β* = 0.6, 95%CI: [0.35, 0.84], p-value = 2.5e-6; post-wake: *β* = 1.4, 95%CI: [1.1, 1.6], p-value = 1e-19; post-awakening power was higher than pre-awakenings (*β* = 1.4 vs 0.6). Disease state (PD/dystonia) showed a statistically significant fixed effect on the rise of subcortical beta power before awakenings (*β* = -0.9, 95%CI: [-1.36, -0.43], p-value = 0.0002) but only a trend after awakenings (post-wake: p-value = 0.06) suggesting a that the rise of subcortical beta is PD specific.

## DISCUSSION

We collected multi-night intracranial brain recordings from four PD and one dystonia participant from cortical and subcortical regions, paired with polysomnography for both DBS ON and OFF conditions, remotely in patients’ own homes over 58 nights. We found increased slow wave activity in the delta band and decreased beta power and connectivity in the cortio-basal network during NREM, an effect that was enhanced by DBS. Within NREM, there was a direct inverse relationship between subcortical beta and cortical delta activity and further, we found that subcortical beta power rises before spontaneous awakenings. These data strengthen the hypothesis that subcortical beta is related to overnight sleep disruptions and spontaneous awakenings in PD.

Our study advances the understanding of sleep neurophysiology in a number of areas. First, technically, we recorded high-resolution intracranial cortical and subcortical neural activity during sleep, over multiple nights (n=58), using a fully embedded, sensing-enabled DBS device in the naturalistic setting from participants in their own homes. Second, we provided evidence of subcortical beta and cortical delta interaction during NREM in PD participants and its modulation by DBS. This effect has been previously noted in a primate model of PD^17^, but to our knowledge, our study demonstrated this interaction in humans with PD for the first time. Third, this is supported by analyses of NREM sleep neurophysiology in both ON and OFF stimulation which disclosed both stimulation-dependent (increased cortical delta and decreased alpha plus low beta in NREM) and stimulation-independent (subcortical beta and cortical delta interaction) effects. Our cortico-basal delta beta interaction finding and pre-awakening subcortical beta rise were significantly stronger effects in the PD cohort in our comparison with our dystonia subject.

It is now established that during the daytime, subcortical beta oscillations are excessive in PD and potentially contribute to circuit disruption and motor symptoms^25,26^. Here, we show that subcortical beta oscillations also disrupt cortical slow oscillations during NREM sleep in humans with PD and are partially responsible for awakenings during the night, validating findings from PD models in primates^17^. Further, we show that DBS stimulation, known to reduce subcortical beta oscillations during wakefulness^27^, here resulted in the increase in cortical delta power and a decrease in cortical alpha and low beta during NREM sleep. This finding aligns with previous studies where an increased accumulation of EEG delta power during NREM sleep was found as a result of subthalamic DBS in PD^28^. One current hypothesis is that DBS therapy improves subjective sleep by reducing overnight discomfort through improved motor movements. Data in our study indicates that DBS therapy appears to additionally improve sleep in PD through direct modulation of beta and delta oscillations.

Although previous studies have documented the presence of subcortical beta oscillations during sleep in STN and GPi^18–22^, these studies to date have been single-night studies and have not probed interactions between beta and delta or spontaneous awakenings during the night nor investigated DBS stimulation effects on sleep physiology. Here, we show that within NREM sleep, subcortical beta inversely correlates with cortical delta power and precedes spontaneous awakenings on a fast time scale. Our findings on the mechanisms of cortical-subcortical interactions during sleep provide a foundation for the development of closed-loop adaptive DBS approaches for restoring normal sleep patterns in people with PD.

The link between sleep dysfunction and daytime motor, mood and cognitive symptoms makes sleep an enticing potential target for further investigation^6–8^. Moreover, sleep disturbances, and particularly reductions in cortical slow wave activity during NREM have been linked to faster disease progression^3,14^. Therefore, targeting beta oscillations during NREM sleep has the potential to reduce overnight insomnia, increase cortical slow waves and improve waking motor and non-motor symptoms. This supports the proposal that daytime neural activities and overnight sleep physiology are notably dissociable and require different strategies for aDBS to optimize rhythms during these two distinct phases. Implementing different aDBS algorithms around the circadian cycle could be achieved by the introduction of daytime (versus sleep) neural classifiers, circadian (clock) based algorithms and combined feedforward and feedback controllers that optimize both daytime and nighttime neurophysiology^29,30^.

Our study has limitations that warrant discussion. First, our ground-truth sleep stage labelings were obtained through a portable polysomnogram and automated sleep-scoring algorithm, validated on healthy controls^31^, instead of a conventional laboratory based PSG. However, we note that our intracranial recordings, grouped according to sleep stages defined from our portable PSG, revealed anticipated and classical changes in ECoG activities across various stages (Supplementary Fig. 4). In particular, the observed elevation in delta power during N3 sleep and reduction in beta power provides evidence of the differentiation of underlying sleep stages within our group of patients using this scheme (Fig 1F). Furthermore, our portable remote setup enabled us to collect multi-night recordings in a natural setting which compares favorably to single-night PSG recordings (from a sleep laboratory) which can be subject to first-night acclimatization effects. We also report a small sample size of participants, although notably, we collected many nights of recordings per subject (n=58 total) which supported highly statistically powered LME analyses that modelled within as well as across subject effects, similar to the strengths of primate research. Our comparison participant was a single cervical dystonia patient (rather than a formal control group) reflecting the uniqueness of this patient cohort, with high-resolution sensing-enabled pulse generators and chronically implanted ECoG electrodes. However, despite this, and in view of the large within-subject dataset size and linear mixed modeling, we were able to show a difference between the dystonia patient and the PD group, which should though be supported in future by larger and more balanced cohorts. Finally, we here restricted our analysis to NREM and canonical power bands with a focus on beta and delta^24^. We did not examine changes in other sleep stages or specifically analyze sleep spindles (which overlap in frequency with low beta) or other frequency bands which will be reported separately.

## CONCLUSIONS

In this study, we recorded and analyzed intracranial FPs with extracranial polysomnography at-home over multiple nights in PD participants, in the presence and absence of DBS. Our data revealed that cortico-basal network power and connectivity in the delta and beta bands are increased and decreased in NREM versus awakening respectively, an effect that was enhanced by DBS. Further, within NREM, cortical delta band slow wave activity was inversely related to subcortical beta, which also rises prior to spontaneous awakenings. These findings uncover a role of subcortical beta in sleep dysfunction in PD and provide targets for future personalized sleep-specific adaptive DBS.

## METHODS

### Participants, demography, and ethics

We recruited 4 participants with idiopathic PD for this study (Table 1). A movement disorders physician diagnosed each individual with PD according to the Movement Disorder Society PD diagnostic criteria^32^. The motor component of the United Parkinson’s Disease Rating Scale (UPDRS) scores were administered by trained raters. We also recruited one participant with cervical dystonia as a comparison subject. Participants were recruited from a parent study focused on investigating closed-loop DBS for daytime motor symptoms. Implanted electrodes were connected to an investigational sensing-enabled Summit RC+S DBS implantable pulse generator provided by Medtronic (Fig 1A)^23^. This study was reviewed by our Institutional Review Board and registered on clinicaltrials.gov (NCT0358289; IDE G180097). The study was also reviewed by the Human Resources Protection Office (HRPO) at Defense Advanced Research Projects Agency (DARPA). Written informed consent was provided by all participants. All subjects had chronic bilateral cortical ECoG electrodes and two PD participants were implanted with bilateral electrodes in the Subthalamic Nucleus (STN; PD2 and PD7) and two PD and 1 dystonia subject were implanted with bilateral electrodes in the Globus Pallidus (GPi; PD3, PD9 and dystonia subject) nuclei (Fig 1B). DBS electrode implantation targets were determined by the clinical team. A movement disorder specialist programmed the patients with conventional DBS settings, optimizing stimulation to address daytime motor symptoms.

### Experimental design and protocols

We collected data using two protocols: long-term multi-night data collection ON stimulation plus separate two night comparison recordings, one night ON DBS and one night OFF DBS. During the long-term overnight data collection, each participant (n=5) was equipped with a portable PSG (Dreem2) headset and overnight intracranial data as well as polysomnography data were recorded for ∼10 nights (Supplementary Table 1) that were predominantly consecutive. In the ON/OFF protocol, overnight data from the PD participants (n=4) were collected for two consecutive days. On the first day DBS was ON (3.075 ± 0.65 mA) and the next day DBS was OFF. During both data collection protocols, the PD participants were on their regular clinical dopaminergic replacement medications. ON/OFF recordings were not completed in the cervical dystonia patient at the patient’s request. All data recordings were performed remotely in patients’ homes.

### Polysomnography acquisition

Extracranial polysomnography (PSG) was recorded through the Dreem headband which includes an automated sleep staging algorithm with extracranial electroencephalography (EEG) data (Dreem2 headband, Dreem Co., Paris, France)^31,33^. The Dreem2 headband provided sleep stage classification hypnograms according to scoring methods (NREM: N3, N2 N1 and REM) of the American Academy of Sleep Medicine (AASM) which has been validated on healthy subjects (Fig 1C)^31,33^. The sleep staging was performed using EEG data at every 30-s epoch. Sleep onset was defined as the start of the NREM sleep (3 consecutive epochs were required to classify N1). Wakefulness after sleep onset (WASO) was calculated as the total waking time after sleep onset and before the last epoch of sleep. As N1 is difficult to detect and physiologically distinct, we focused our analysis on N2 and N3 stages for NREM sleep.

### Intracranial data collection

For each participant, the Summit RC+S device was implanted bilaterally and connected to bilateral sensing and stimulation-capable quadripolar leads in the basal ganglia targets (STN in 2 PD patients or GPi in 2 PD patients and 1 cervical dystonia patient) plus quadripolar sensorimotor chronic electrocorticography (ECoG), sensing only strips, with 4 electrode contacts spanning the central gyrus (Fig 1B). Overnight intracranial data were collected from cortical and subcortical structures in both left and right hemispheres (Fig 1D) in addition to data from bilateral accelerometers embedded within the chest mounted pulse generator devices. The time series FP data were recorded at either a 250 Hz or 500 Hz sampling rate.

### Data Preprocessing

The intracranial recordings were validated and synchronized to the PSG recordings using accelerometry data. Cross-correlation was applied to accelerometry data from both the Dreem2 band and the RC+S neurostimulator in order to ascertain the delay between PSG and RC+S time series (Supplementary Fig. 3). As PSG hypnogram sleep stage estimates were performed on 30-s epochs, we also used post waking movement (measured via accelerometry) to further re-align to waking at a sub 30-s scale (Supplementary Fig. 3). All intracranial data were downsampled to 250 Hz and filtered through a 0.8-100 Hz zero-phase IIR elliptic bandpass filter with 1dB passband ripple and 100 dB attenuation (‘filtfilt’ and ‘designfilt’ function in Matlab). Large artifactual spikes in the subcortical intracranial data were removed along with the corresponding cortical data (Supplementary Fig.3). To identify artifacts, absolute squared subcortical data were first smoothed with a Gaussian kernel with 1s window then any period larger than 5 times the median over the whole night was considered artifactual spikes. The ECG artifacts in the subcortical data were removed using a combination of two ECG data remover algorithms (‘PerceptHammer’ and ‘Perceive’ library; Matlab; Supplementary Fig. 3)^34,35^.

### Power spectrum analysis

To calculate the power spectra, the intracranial data from each night were z-scored for each location. Then, the NREM data segments (N2+N3) were collected together according to the PSG hypnogram labels. The selected data were segmented into 5-s epochs and power spectra were calculated for each epoch using a Hamming window of 1-s, 512 point FFT with 50% overlap by Welch’s method (‘pwelch’ in Matlab) which was normalized by the total power in 0-50 Hz. The calculated power spectrums for each epoch were then pooled over both hemispheres within subjects. For calculating the change in power spectrum in NREM with wake as the baseline, the power spectrum for wake epochs were calculated in a similar manner as during NREM and the difference between the average wake power spectrum and NREM power spectrum for each night was calculated. For calculating the ON vs OFF power spectrum, average power spectra were calculated for ON and OFF nights and their difference was taken. The averages were calculated on log-transformed power spectra.

### Spectral coherence analysis

To compute the spectral coherence, the intracranial data obtained from each night were normalized using z-scoring for each location. Subsequently, NREM data segments comprising N2 and N3 sleep stages were extracted and then divided into 5-s epochs. For each epoch, 5-s of cortical and subcortical data were utilized for estimating the one-sided magnitude squared coherence using the multitaper method (‘mscohere’; Matlab) with Hamming window of 1-s and 512 point FFT. The epoch-wise spectral coherences were then pooled over both hemispheres. Similarly to the power spectral analysis, spectral coherence for wake epochs were calculated and the difference between the average of wake and NREM spectral coherence for each night was calculated in order to obtain the change in spectral coherence in NREM with wake as the baseline.

### Beta-delta correlation analysis

To analyze the interaction between subcortical beta and cortical delta activity during NREM sleep in intracranial signals, we applied z-scoring, power spectrum calculation and normalization techniques as previously described. However, there was one exception regarding the normalization of the cortical power spectrum where instead of normalizing it by dividing the total power (0-50 Hz), we divided it by the total power excluding the beta range (0-13 and 31-50 Hz). This adjustment was necessary to avoid detecting spurious negative correlations that could be caused by the normalization procedure itself. Both subcortical beta and cortical delta were calculated for 5-s epochs which were log-transformed for each night and each hemisphere. The band powers were then pooled over both hemispheres. Subsequently, for each participant, we calculated Spearman’s rho correlation coefficient between subcortical beta and cortical delta power across all 5-s epochs for each night. Similar results were obtained when employing various other normalization methods. For calculating the delay between subcortical beta and cortical delta power, normalized cross-correlation (‘xcorr’ function in Matlab) was calculated between these band powers from the 5s epochs from above for each night. Lag was calculated by finding the minimum (trough, reflecting a negative relationship) normalized cross-correlation between the two band powers. Epoch band powers for each night were smoothed using a 20-point Gaussian kernel. Data for each night were mean-subtracted and pooled from both hemispheres. To investigate interactions between delta and beta powers from cortex, we applied the same power spectrum calculation techniques on 5s epochs as previously described in beta-delta correlation analyses. The only exception was the normalization step of the power spectrum which was not applied to avoid detecting artificial negative correlations that could be imposed by the normalization of the power spectrum. Spearman’s rho correlation coefficient was calculated between the cortical delta and beta power in all 5s epochs throughout each night for all participants.

### NREM to wake transition analysis

To investigate the changes in spectral power during NREM to wake events, all intracranial data were bandpassed using the zero-phase IIR elliptic bandpass filters. Next, the data were z-scored for each night for each hemisphere at all locations. Hilbert transform was applied to the z-scored data and the absolute square of the results were converted into a decibel scale for band-specific power. After detecting all NREM to wake transitions, events with NREM sleep less than 85-s and Wake period of less than 25-s were ignored. Maximum 1-s total discontinuity in the sleep event was allowed and band-power for each 5-s epoch was averaged. All epochs in NREM data after 40-s from NREM onset and 40-s before awakening were averaged to calculate deep NREM (slow-wave sleep; SWS) power. All epochs in awake stage data after 25-s from wake onset were averaged to calculate awake stage power. All data were analyzed from the ON stimulation multi-night dataset.

### Statistical methods

A significance threshold of 0.05 was employed to determine statistical significance. Linear mixed effect models (LME) were utilized (‘fitlme’ in Matlab) for investigating the spectral power and coherence differences, the interactions between cortical and subcortical beta with cortical delta powers. Theoretical likelihood ratio test (‘compare’ in Matlab) was used for comparing LME models. All analyses were performed using Matlab 2022a (Mathworks).

## DATA AVAILABILITY

Datasets and codes generated and/or analyzed in this study will be shared upon reasonable request.

## ACKNOWLEDGMENTS

This work was funded by the Defense Advanced Research Projects Agency (DARPA) under grant HR0011-20-2-0028 and the National Institute of Neurological Disorders and Stroke (NINDS) under grant 1R01NS131405-01. Google research provided unrestricted laboratory gift support for our research. We express our gratitude to the participants for their valuable contributions to this study. DARPA, the NINDS and Google research played no role in the study design, data collection, analysis, and interpretation of data, or the writing of this manuscript. We thank and acknowledge Shravanan Ravi for his help supporting data collection and Tom Wozny for his contributions in visualizing ECoG and DBS lead locations in Figure 1B. Thanks to all the patients who joined the study and to the staff in the Department of Neurology of the University of California San Francisco for their help and support.

