## Supplementary for "Multi-night naturalistic cortico-basal recordings reveal mechanisms of NREM slow wave suppression and spontaneous awakenings in Parkinson’s disease"

**Supplementary Table 1: Sleep statistics**

| | ON stimulation | | | | | OFF stimulation | | | |
|---|---|---|---|---|---|---|---|---|---|
| | DYS | PD3 | PD9 | PD2 | PD7 | PD3 | PD9 | PD2 | PD7 |
| SO (min) | 18.8±2.4 | 29.9±3.7 | 32.2±3.4 | 19.2±1.9 | 25.3±2.9 | 22.4 | 106.4 | 34.9 | 22.5 |
| N1 (min) | 23.5±2.7 | 33.5±2.7 | 28.9±2.3 | 35.7±3.0 | 39.8±1.3 | 17.5 | 33 | 47.9 | 25 |
| N2 (min) | 143.0±13.0 | 99.2±9.9 | 184.6±11.3 | 181.8±10.0 | 192.4±14.9 | 92.8 | 169 | 236.9 | 232.6 |
| N3 (min) | 99.5±6.1 | 204.9±10.2 | 35.8±4.8 | 54.5±5.8 | 71.4±3.9 | 150.1 | 22.5 | 75.8 | 60.9 |
| REM (min) | 55.8±6.3 | 112.8±8.0 | 125.6±14.8 | 59.5±5.4 | 93.2±13.1 | 140.6 | 72.5 | 62.8 | 92.8 |
| N2+N3 (min) | 242.5±12.9 | 304.2±14.4 | 220.4±12.1 | 236.3±9.1 | 263.8±14.6 | 242.9 | 191.5 | 312.8 | 293.5 |
| WASO (min) | 62.0±12.2 | 73.5±8.2 | 17.7±3.2 | 63.1±9.2 | 39.0±4.8 | 132.7 | 67.5 | 102.2 | 28.9 |
| Wake event | 24.7±2.5 | 36.4±2.0 | 16.5±2.2 | 20.7±2.4 | 38.8±3.4 | 21 | 14 | 35 | 23 |
| TST (hours) | 6.4±0.2 | 8.7±0.1 | 6.5±0.2 | 6.6±0.3 | 7.3±0.3 | 8.9 | 6.1 | 8.8 | 7.3 |
| Total nights | 9 | 11 | 12 | 12 | 10 | 1 | 1 | 1 | 1 |

SO = Time to Sleep onset; WASO= Wake after sleep onset; N1, N2, N3, REM, N2+N3 total duration times per night in minutes; TST= total sleep time; Wake event = total wake events during one night.

ON stimulation includes average sleep metrics for 11 nights of recordings at home. OFF stimulation includes a single night of at home recording in the absence of stimulation.

**Supplementary Table 2: Performance of wake prediction**

| | Deep vs pre-wake (-5s) NREM | | | | | Deep NREM vs post-wake (+15s) | | | | |
|---|---|---|---|---|---|---|---|---|---|---|
| Subject | Dys | PD3 | PD9 | PD2 | PD7 | Dys | PD3 | PD9 | PD2 | PD7 |
| Accuracy | 63.7 | 73.7 | 55.6 | 68.9 | 69.8 | 65.3 | 78.9 | 69.4 | 84 | 77.4 |
| AUC | 63.3 | 73.4 | 59 | 77.4 | 71.3 | 60.6 | 83.9 | 74.8 | 94.9 | 80.7 |
| Sensitivity | 33.9 | 63.2 | 36.1 | 41.5 | 50.9 | 37.1 | 73.7 | 63.9 | 71.7 | 66 |
| Specificity | 93.5 | 84.2 | 75 | 96.2 | 88.7 | 93.5 | 84.2 | 75 | 96.2 | 88.7 |
| PPV | 84 | 80 | 59.1 | 91.7 | 81.8 | 85.2 | 82.3 | 71.9 | 95 | 85.4 |
| NPV | 58.6 | 69.6 | 54 | 62.2 | 64.4 | 59.8 | 76.2 | 67.5 | 77.3 | 72.3 |
| Odds ratio | 7.4 | 9.1 | 1.7 | 18.1 | 8.1 | 8.5 | 14.9 | 5.3 | 64.6 | 15.2 |
| U-test p-value | 0.01 | 0.01 | 0.2 | 1.2e-6 | 1.6e-4 | 0.04 | 3.7e-4 | 3e-4 | 1e-15 | 5e-8 |

Individual QDA model performance for binary classification. PPV = positive predictive value, NPV = negative predictive value, U-test = Wilcoxon rank sum test, AUC = Area under the receiver operating characteristic curve

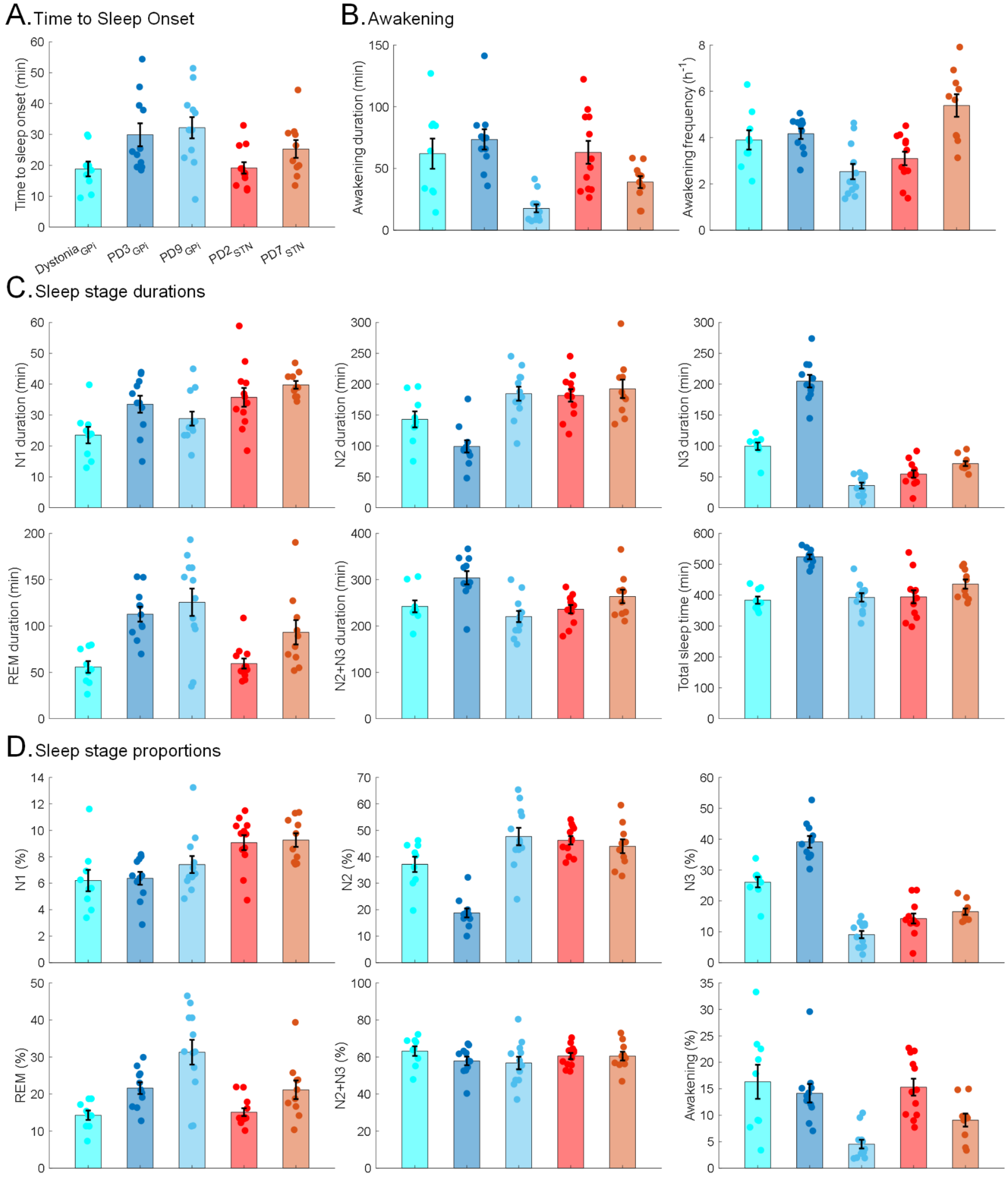


**Supplemental Figure 1: Sleep statistics in ON stimulation.** Sleep statistics (mean ± SEM ) for all participants (n=5) during overnight recordings with ON stimulation. **(A)** Time to sleep onset **(B)** Wake after sleep onset (awakening during the sleep) in total duration and frequency over one night **(C)** Durations of all sleep stages in minutes **(D)** Total proportion of sleep stages.

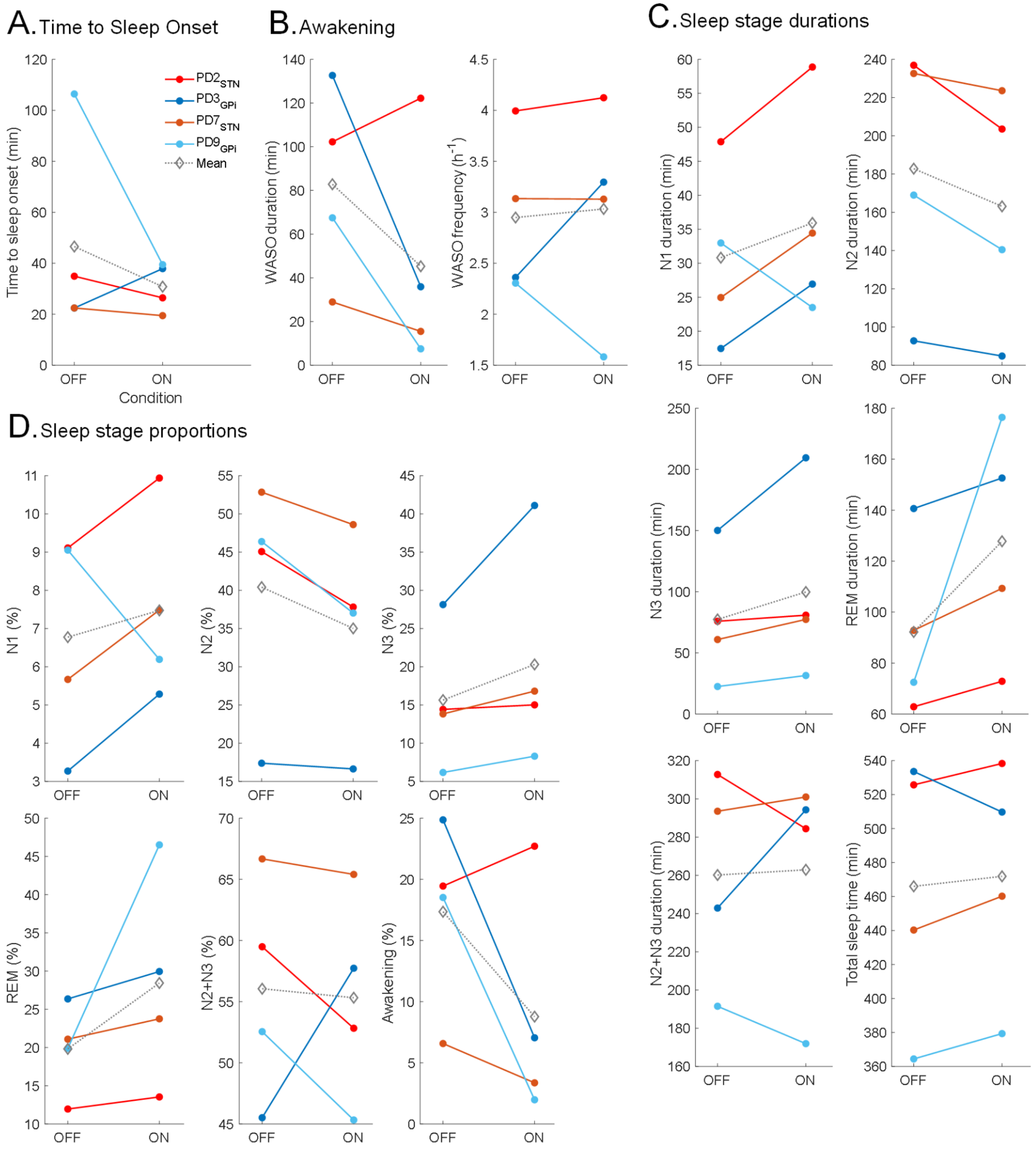


**Supplemental Figure 2: ON vs OFF stimulation sleep statistics.** Sleep statistics for all PD participants (n=4) during overnight recordings during consecutive one night ON and one night OFF stimulation conditions. **(A)** Time to sleep onset **(B)** Wake after sleep onset (awakening during the sleep) in total duration and frequency over one night **(C)** Durations of all sleep stages in minutes **(D)** Total proportion of sleep stages. x-axes are stimulation conditions (ON/OFF) and the dashed gray lines show average across all PD participants.

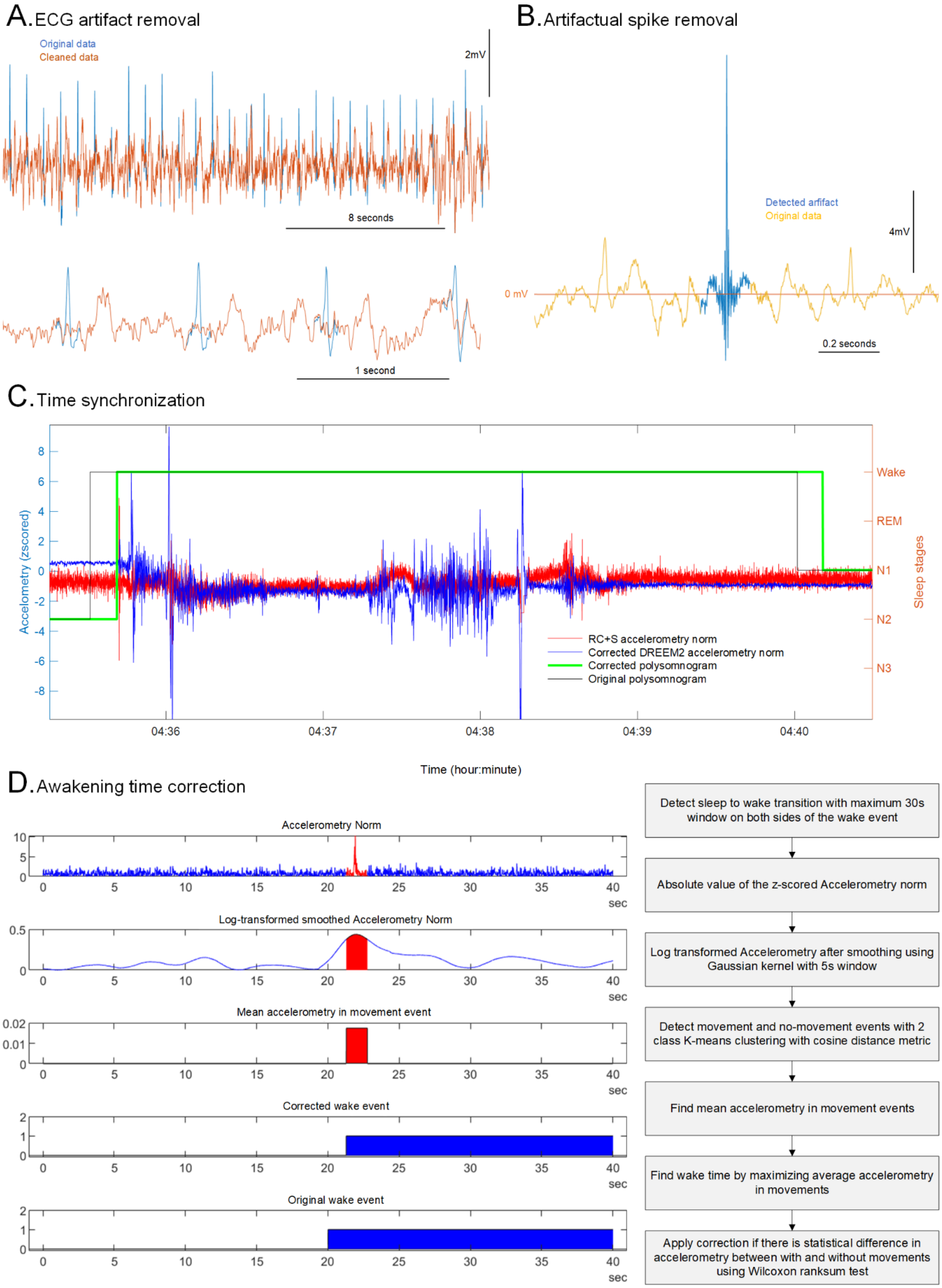
A. ECG artifact removal
Original data
Cleaned data
2mV
8 seconds
1 second
B. Artifactual spike removal
Detected arifact
Original data
4mV
0 mV
0.2 seconds
C. Time synchronization
Accelometry (zscored)
Sleep stages
Wake
REM
N1
N2
N3
RC+S accelerometry norm
Corrected DREEM2 accelerometry norm
Corrected polysomnogram
Original polysomnogram
04:36
04:37
04:38
04:39
04:40
Time (hour:minute)
D. Awakening time correction
Accelerometry Norm
Log-transformed smoothed Accelerometry Norm
Mean accelerometry in movement event
Corrected wake event
Original wake event
sec
Detect sleep to wake transition with maximum 30s window on both sides of the wake event
Absolute value of the z-scored Accelerometry norm
Log transformed Accelerometry after smoothing using Gaussian kernel with 5s window
Detect movement and no-movement events with 2 class K-means clustering with cosine distance metric
Find mean accelerometry in movement events
Find wake time by maximizing average accelerometry in movements
Apply correction if there is statistical difference in accelerometry between with and without movements using Wilcoxon ranksum test

**Supplemental Figure 3: Data processing procedures.** Removal of the ECG artifact **(A)** and movement-related spike artifact **(B)** from the RC+S field potential data. **(C)** Time synchronization of polysomnogram from DREEM2 with intracranial data streams from RC+S devices using accelerometry data as references **(D)** Awakening time correction using accelerometry data and by finding movement events.

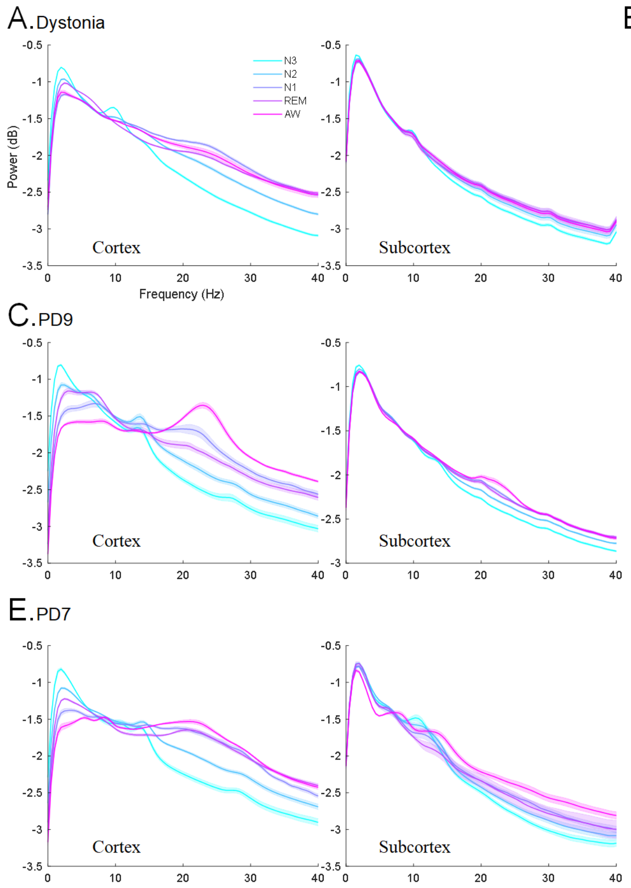


**Supplemental Figure 4.** Spectral power (mean ± SEM ) in cortex and subcortex for all sleep stages in each participant (n=5). Data from ON stimulation. Data is averaged over 10 nights and bilateral hemispheres.
